# Transcriptomic and functional analysis of the oosome, a unique form of germ plasm in the wasp *Nasonia vitripennis*

**DOI:** 10.1101/384032

**Authors:** Honghu Quan, Jeremy Lynch

## Abstract

**Background:** The oosome is the germline determinant in the wasp *Nasonia vitripennis* and is homologous to the polar granules of *Drosophila*. Despite a common evolutionary origin and developmental role, the oosome is morphologically quite distinct from polar granules. It is a solid sphere that migrates within the cytoplasm before budding out and forming pole cells.

**Results:** To gain an understanding of both the molecular basis of the novel form of the oosome, and the conserved essential features of germ plasm, we quantified and compared transcript levels between embryo fragments that contained the oosome, and those that did not. The identity of the localized transcripts indicated that *Nasonia* uses different molecules to carry out conserved germ plasm functions. In addition, functional testing of a sample of localized transcripts revealed potentially novel mechanisms of ribonucleoprotein assembly and pole cell cellularization in the wasp.

**Conclusions:** Our results demonstrate that numerous novel and unexpected molecules have been recruited in order produce the unique characteristics of the oosome and pole cell formation in *Nasonia*. This work will serve as the basis for further investigation into the patterns of germline determinant evolution among insects, the molecular basis of extreme morphology of ribonucleoproteins, and the incorporation of novel components into developmental networks.

## Background

Germline establishment is a crucial event for sexually reproducing organisms. Germline cells are special in that they are able to generate all of the cell fates of the soma and to regenerate themselves. There are two major strategies to specify the germline among animals: zygotic induction and maternal provision. In zygotic induction, inductive signals from surrounding tissues drive the establishment of germline fate, usually relatively late in embryogenesis, after the transition from maternal to zygotic control [1]. In contrast, in the maternal provision mode, the germ cells are specified by determinants called germ plasm that are synthesized and localized during oogenesis and are the first group cells formed during embryogenesis. Germline specification in this mode occurs very early in development, usually prior to the activation of the zygotic genome [1, 2]. Classical experiments have shown that germ plasm is both necessary and sufficient to establish the germline fate [3-5].

It is likely that the maternal provision mode of germ plasm evolved multiple times among the animals, and this reflected in the molecular basis of germ plasm determinants [6-27]. In vertebrates, germ plasm (where it exists) is dependent on the maternal localization of *bucky ball* [28]. In insects, the gene products of *oskar* (*osk*) are both necessary and sufficient to induce germ plasm, and thus, primordial germ cells (PGCs) [29-32]. Downstream of these nucleators is a suite of highly conserved germline-associated molecules (i.e., Vasa (Vas), Nanos (Nos), Tudor (Tud), etc.) that are recruited to the germ plasm where they presumably carry out functions ancestral for the animal germline [30, 33-35].

There are several conserved properties of PGCs that must be conferred to naive cells by germ plasm. One of these is a period of transcriptional quiescence germ cells undergo after being specified. This feature makes the composition of a germ plasm even more important since it means that mRNAs and proteins critical for at least the early germ cell functions need to be provided in the germ plasm itself. Germ cells also usually become highly migratory later in development as they seek to colonize the developing gonad. They are often highly enriched for mitochondria and have specific metabolic needs [36]. Since they carry the genome that will be passed to future generations, germ cells have enhanced mechanisms to prevent DNA damage and to reduce the activity of transposable elements [37, 38]. Finally, they must have the capability to induce pluripotency to their genome, as germline cells will be the source for all cell fates in the eventual progeny.

Like all developmental processes, there is likely to be variation in the details of these conserved features of germline determination, whether due to selective or neutral forces. Pressures that could impact the composition of the germ plasm could be differential activity of transposable elements in the genome, or a novel path for migration of the PGCs to the gonad, for example. Novelties in embryogenesis may also drive germ plasm composition. For example, in the parthogenetic, paedomorphic embryos of the midge *Miastor,* somatic nuclei undergo significant chromosomal diminution in the early cleavages while the cell that takes up germ plasm maintains the full complement of chromosomes, indicating that the germ plasm contains a component that prevents the loss of chromosomes during the early cleavages [39].

Holometabolous insects are an ideal system with which to study how germ plasm evolves, given that it is ancestral in this clade [35], the unparalleled levels of diversity, and the strong baseline understanding germ plasm function derived from *Drosophila melanogaster*. Here we focus on the parasitic wasp *Nasonia vitripennis* as a model to compare to the fruit fly. Like *Drosophila, Nasonia* depends on Osk, Vas, and Tud to assemble germ plasm [35, 40]. However, in contrast to the collection of small granules stably associated with the posterior pole that make up the *Drosophila* germ plasm, the *Nasonia* germ plasm forms a very large, dense organelle called the oosome (Figure 1). This highly divergent morphology strongly indicates that the composition of the *Nasonia* oosome may diverge significantly from the polar granules of *Drosophila*.

**Fig. 1.**
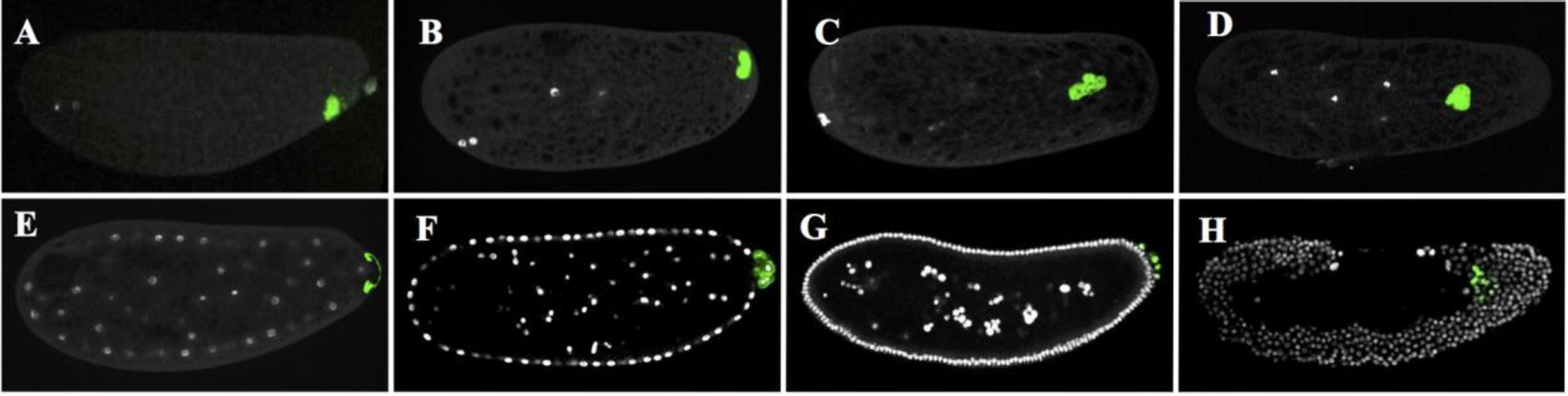
The complete process of *Nasonia* germline development marked by the novel gene *Nv-bark*. A-D: the green shows the oosome shapes and localizations in different nuclear divisions during pre-blastoderm stage. E: the oosome flattenes into the bud (green) at the beginning of blastoderm stage. F: the first batch of pole cells (green) is formed from the bud during early syncytial blastoderm stage. G: the pole cells divide several times to make more and smaller cells (green) during cellularized blastoderm stage. H: the pole cells (green) has entered into the embryo and migrate to the gonads during gastrulation stage. The embryos are arranged according to their embryogenesis stages with the posterior side to right and dorsal side on the top. DAPI (white) marks the nuclei.

The behaviors of the oosome and the PGCs in *Nasonia* further imply a divergent composition of the oosome. In freshly laid eggs, the oosome is tightly bound to the ventral-posterior cortex of the embryo (Fig. 1A). When the zygotic nucleus forms and moves into the interior of the embryo, the oosome detaches from the cortex and coalesces into a dense, extremely large structure in the same central column as the nuclei. It migrates anteriorly, sometimes being found near to 50% egg length, before migrating back to the posterior pole (Fig. 1B-D). As the cleavage nuclei migrate toward the cortex, the oosome flattens into a crescent on the posterior pole of the embryo while a large bud protrudes from the pole (Fig. 1E). Typically, two or three nuclei become associated with the bud and the oosome material. The bud pinches off, and the nuclei rapidly individuate into pole cells (Fig. 1F,G).

This is distinct from pole cell formation in *Drosophila,* where each pole cell forms individually upon association with a certain critical amount of pole plasm [41]. After dividing a few times, the pole cells remain stable at the posterior pole of the egg until gastrulation, when they migrate through the posterior epithelium and coalesce into to two masses, presumably where the primordial gonads are developing (Fig. 1H). This migration is distinct from *Drosophila* pole cells which remain external on the posterior pole as the germ band undergoes massive extension well after gastrulation and migrate internally through the gut wall [41].

Thus, it is clear that *Nasonia* and *Drosophila* share some fundamental aspects of germline establishment, but they also have their own diverged features that fit in their own embryogenesis program. This raises the question of which genes are the core components for the maternal provision mode and which genes contribute to their own distinct features in germline development.

To address these questions, we compared the mRNA content of anterior and posterior halves of the pre-blastoderm stage *Nasonia* embryos in an effort to identify the components specifically localized to the oosome. We found only a few of mRNAs conserved in both fly polar granules and the *Nasonia* oosome, such as *osk, nos*, and *ovo* (see Supplemental File 1). The rest are all novel components germ plasm that either lack *Drosophila* homologs or have homologs in *Drosophila* that do not play any roles in germline development. We performed functional studies for a set of localized transcripts, all of which showed roles in the unique features of *Nasonia* germline development, demonstrating the value of our approach to identify the molecular sources of novelty among various insect lineages.

## Results

### RNA-seq analyses of the anterior and posterior poles of the wasp *Nasonia* early embryos

To identify the maternal transcripts in the oosome, we isolated the total RNA separately from anterior and posterior poles of the pre-blastoderm stage *Nasonia vitripennis* embryos. Six samples that comprised three sets of each pole were sequenced and subjected to differential expression analysis, which are described in more detail in the Methods.

Our first attempt yielded encouraging, but mixed results. Several novel posteriorly enriched factors were identified, and we did recover some known posterior factors (*Nv-osk* and *Nv-nos*). However, other known localized factors such as *Nv-cad* and *Nv-dpp* were not found to be statistically significant. In addition, one of the posterior samples was of poor quality, and could not be used in the differential expression analyses, reducing the power of our approach. Since we wanted results as comprehensive as possible, we repeated the experiment using a few adjustments (see Methods) and obtained much more robust results. While this second experiment was being prepared, we moved forward with expression and functional analysis of novel candidates identified in the first. Since a handful of transcripts with confirmed oosome localization were found only in the first experiment, the remainder of the results will include these transcripts along with those identified in the second experiment.

All known maternally localized molecules (*Nv-osk, nos, dpp, cad, otd1*, and *gt*) were found with high significance, giving confidence that our experimental design and analysis approaches were appropriate for our goal of discovering the mRNAs localized to the oosome. Overall, we found 92 transcripts with apparent significant enrichment at the posterior pole. These ranged in levels of enrichment from 1.4 to 55 times higher in the posterior fragments compared to anterior fragments. Our analyses also uncovered anteriorly enriched mRNAs, of which there were 89, with a range of fold enrichment from 1.4 to 10 times higher at the anterior.

### Novel transcripts localized in the anterior half of the *Nasonia* early embryos

While the anterior factors are not the focus of this manuscript, some interesting observations were made in examining a small sample of candidate mRNAs. Most transcripts are localized in small domains at the anterior cortex and seem to extend toward the posterior in variable tendrils, rather than being uniform or graded caps of anterior localization (Fig. 2A1-K1). A notable exception is the transcript of *Nasonia* homolog of *mex-3,* an RNA binding protein known for controlling translation of orthologs of the posterior patterning factor *caudal* in the nematode *C. elegans* and the beetle *Tribolium* [24, 42]. *Nv-mex3* mRNA is localized in a broad domain extending far toward the posterior of the embryo (Fig. 2L1). *Nv-mex3* expression is highly dynamic and variable in both the blastoderm and post-gastrular stages (Fig. 2L2-L3, additional images not shown).

**Fig. 2.**
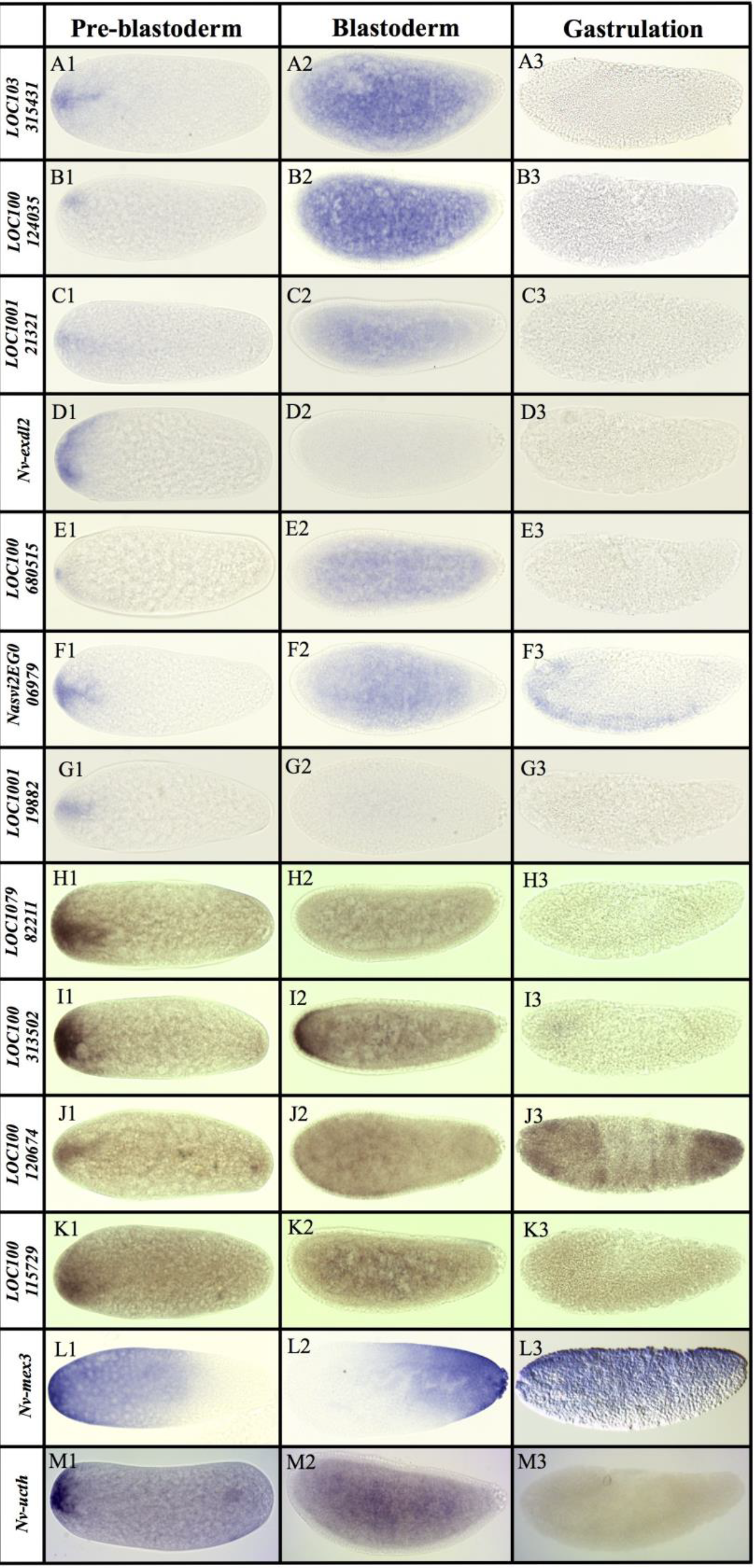
Genes expressed anteriorly in the pre-blastoderm stage. All embryos are aligned and grouped into three columns (Pre-blastoderm, Blastoderm and Gastrulation) according to their embryogenesis stages, with posterior side to the right and dorsal side on the top.

Transient localization is the most common feature of the anteriorly localized transcripts. Most are ubiquitous or absent by the time the early syncytial blastoderm forms (Fig. 2 A2-H2, J1-K3, M1-M3), except for *LOC100313502* which persists into the blastoderm stage where it forms a anterior cap (Fig. 2 I1-I3). Among the anteriorly enriched transcripts, many have predicted functions that may be relevant to egg activation (e.g. four encoding ion channels), and anterior-posterior polarity and patterning (e.g. transcription and translation factors, see Supplemental file 1). There are also a large number of transcripts with no clear homologs outside of *Nasonia* see (Additional File 1). One of these (*LOC100119982*, Fig. 2G1-G3) is a member of a novel family of ankyrin domain containing molecules that are specific to Chalcid wasps and appear to have obtained a broad diversity of expression and potential function during *Nasonia* development [43].

Finally, a handful of molecules localized at both poles were detected. These include the known factors *Nv-otd1* [44] and *Nv-TollC* (JAL in preparation), along with *Nv-endoglucanase* (Fig. 3E), *Nv-insulin-like growth factor (Nv-igf,* Fig. 3C*), Nv-ucth* (a ubiquitin carboxy terminus hydrolase, Fig. 2M)). The fact that these were found despite lower apparent fold differences between the two embryonic halves, further gave confidence that our analysis was robust enough to detect even subtle germ plasm localization of the vast majority of mRNAs.

**Fig. 3.**
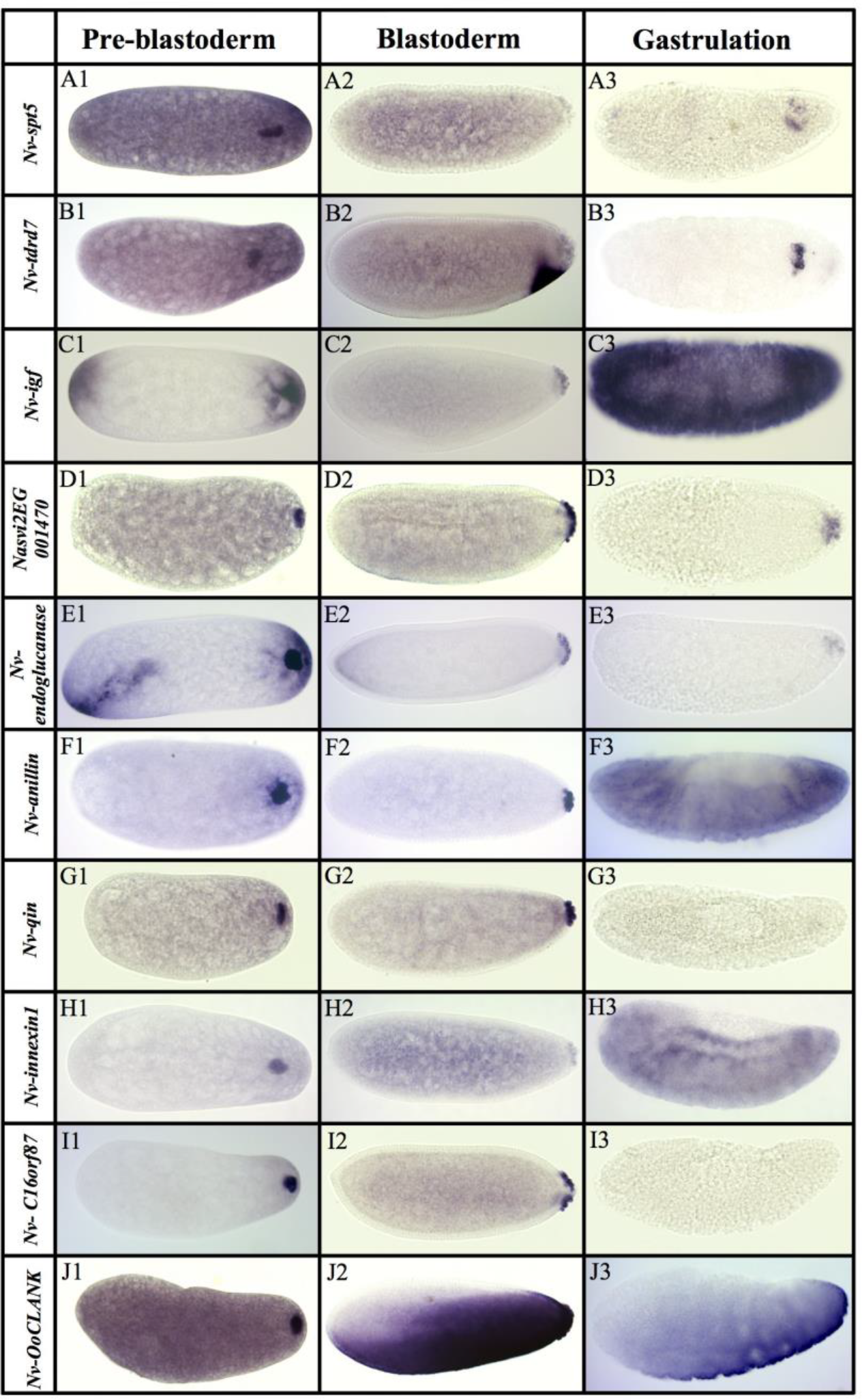
Transcripts localized to the oosome that are subsequently maintained in pole cells. All embryos are aligned and grouped into three columns (Pre-blastoderm, Blastoderm and Gastrulation) according to their embryogenesis stages, with posterior side to the right and dorsal side on the top.

### General description of the novel transcripts localized in the posterior pole of the *Nasonia* early embryos

From the two analyses, we identified 92 candidate transcripts that were statistically enriched at the posterior half of the *Nasonia* embryos. We then isolated PCR products for 54 of these genes and made probes to determine their expression patterns during early embryogenesis. We were able to confirm 47 transcripts that are expressed posteriorly during pre-blastoderm stage. Of the remaining transcripts, one of them was not successfully cloned, and the rest were successfully cloned but did not have any specific, localized expression. Interpretation of these potential false positives (as well as some false negatives) will be discussed later. Among the 28 transcripts with expression in the posterior half of the embryo, we grouped them into three categories based on their expression patterns: 1) genes expressed in oosome and with expression maintained in the PGCs (Fig. 3), 2) mRNAs localized in the oosome but then degraded in the PGCs (Fig. 4), and 3) mRNAs strongly enriched in the posterior region of the embryo, but not incorporated in the oosome (Fig. 5).

**Fig. 4.**
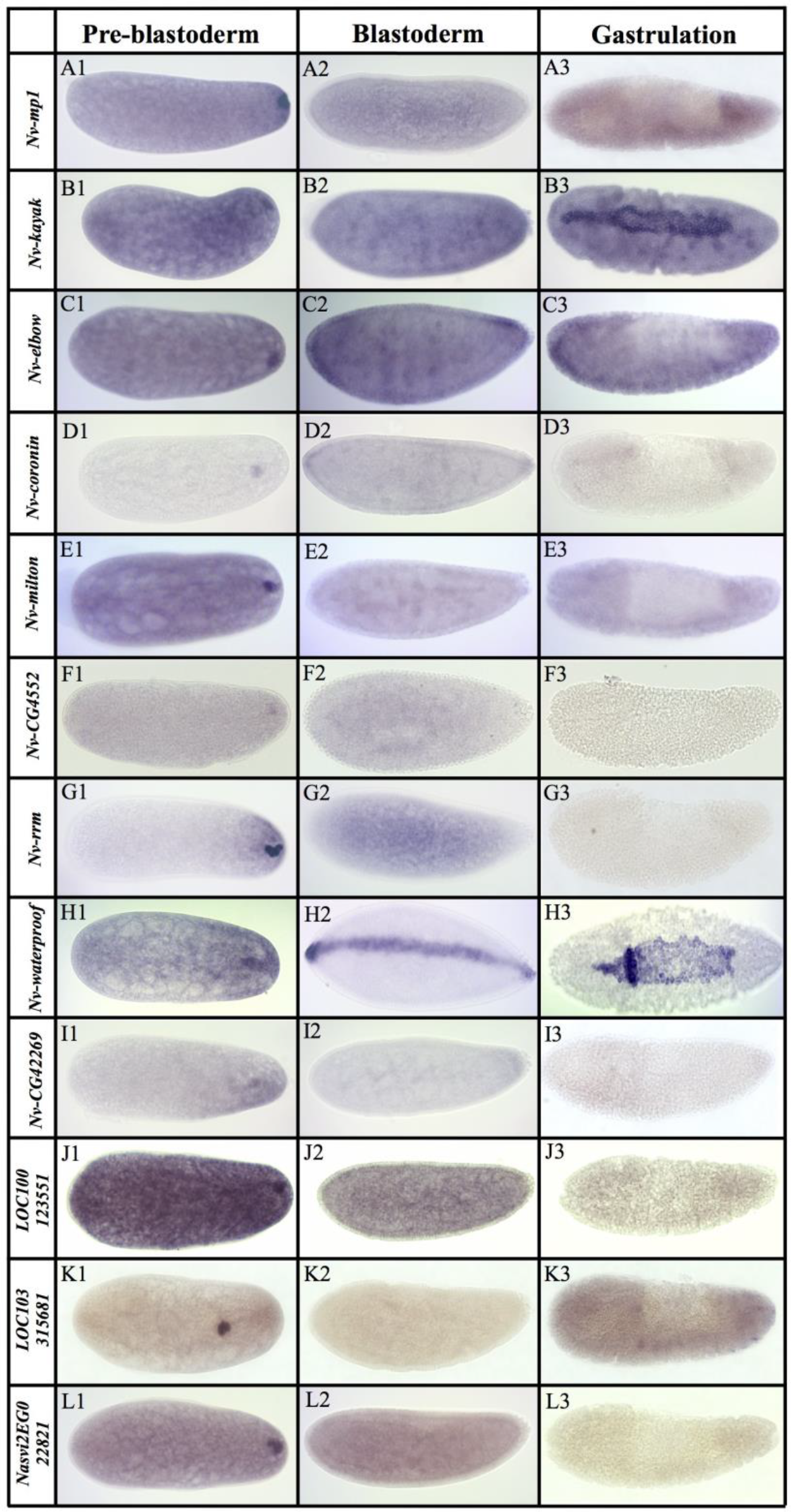
mRNAs localized in the oosome (Pre-blastoderm) but are excluded from the pole cells. All embryos are aligned and grouped into three columns (Pre-blastoderm, Blastoderm and Gastrulation) according to their embryogenesis stages, with posterior side to the right and dorsal side on the top.

**Fig. 5.**
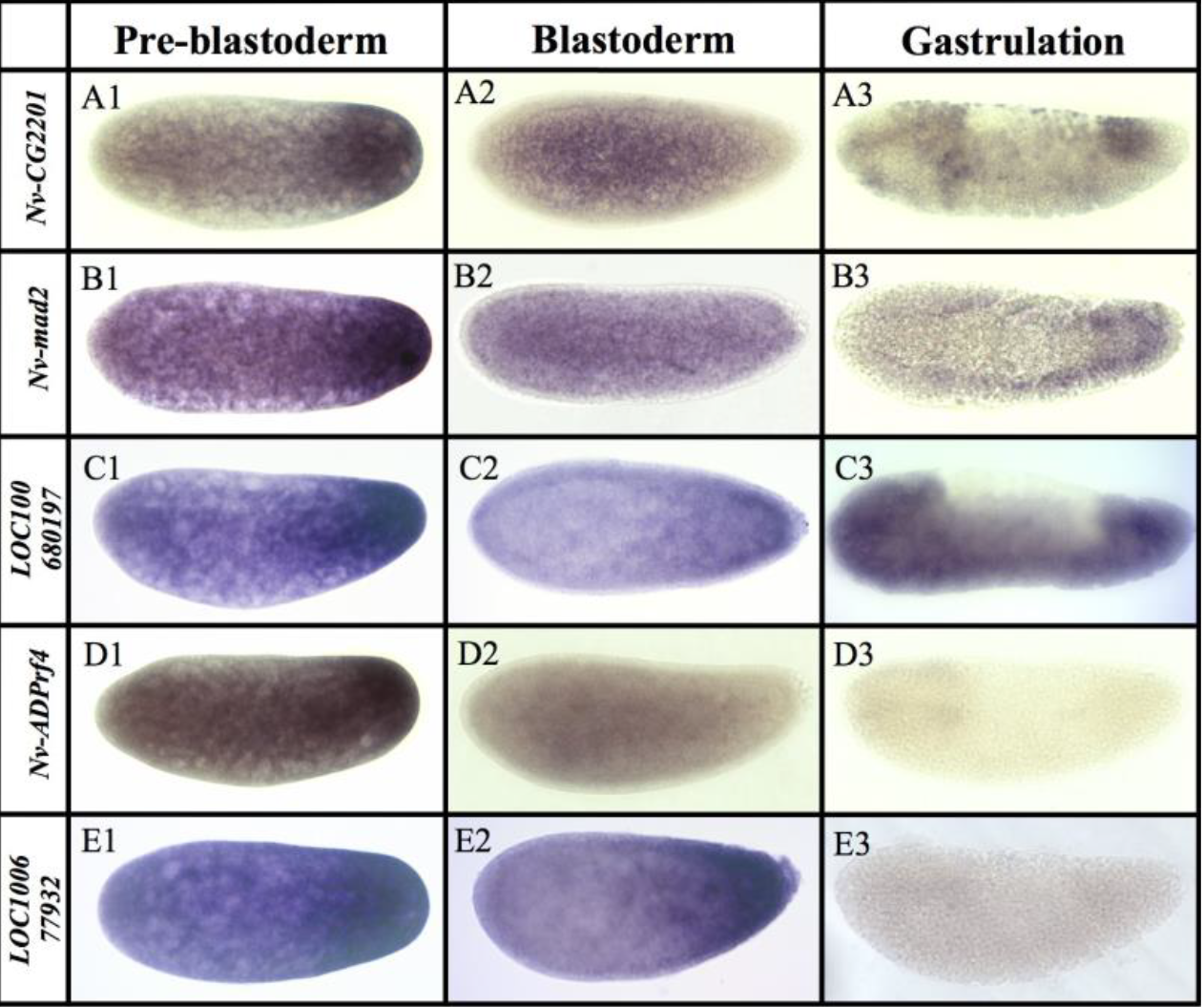
mRNAs strongly enriched in the posterior region of the embryos, but not enriched in the oosome. All embryos are aligned and grouped into three columns (Pre-blastoderm, Blastoderm and Gastrulation) according to their embryogenesis stages, with posterior side to the right and dorsal side on the top.

### Highly germline-associated transcripts

We identified 11 transcripts that are localized to the oosome and maintained in the PGCs at pole cell formation and beyond. We consider these to be the strongest candidates for having important roles in the specification and function of the PGCs but cannot exclude *a priori* that they have other (or no) important functions. Six transcripts are maintained throughout the stages of germline development followed in this manuscript: oosome, pole cells, and migrating germ cells. They include *Nasonia* homologs of the fly genes *bark beetle* (*Nv-bark*) (Fig. 1), *spt5* (*Nv-spt5*), *tejas* (*Nv-tdrd7*), *insulin-like growth factor* (*Nv-igf*), and two transcripts without fly homologs (*Nasvi2EG001470* and *Nv-endoglucanase*) (Fig. 3A1-E3).

Among these, *Nv-bark* is the best germline marker, bearing strong and consistent germline association throughout early development, including expression in the late embryonic gonads (Fig. 1). In this respect, it is better than our previously favored marker, *Nv-nos*, which is downregulated significantly toward the end of PGC migration [40]. Bark is a large transmembrane protein and is not expressed in the *Drosophila* germline. Its only known role is in stabilizing tricellular junction in epithelial cells during embryogenesis [45, 46]. It is not clear how this function is relevant to germline function, and indicates a novel recruitment of this factor in the wasp.

*Nv-spt5* is significantly enriched in the oosome with low levels of ubiquitous expression in the rest of the embryos in pre-blastoderm stage (Fig. 3A1). In the blastoderm and migrating germ cell stages expression is reduced but still enriched in the pole cells while the ubiquitous expression in the embryo persists (Fig. 3A2-A3). Spt5 homologs are involved in regulating RNA polymerase progression during transcription [47], which might indicate that Nv-Spt5 is involved in repressing or otherwise regulating the onset of transcription in the germ cells.

*Nv-tdrd7* is present at appreciable levels throughout the bulk cytoplasm and is also strongly localized in the oosome (Fig. 3B1). This pattern is well reflected in the quantification of mRNA levels in the two halves of the embryo, which show significant numbers of reads coming from the anterior half of the embryo. At the blastoderm stage, *Nv-tdrd7* is moderately enriched in the pole cells and is zygotically expressed in a ventral-posterior patch (Fig. 3B2), which was detected in our earlier analysis of dorsal-ventral patterning [48]. After gastrulation, *Nv-tdrd7* is strongly upregulated in a group of cells that are near to where the germ cells migrate, but it is not clear if they are germ cells (Fig. 3B3).

*Nv-igf* is initially expressed in a bipolar pattern (similar to *Nv-otd1*[44]), before becoming specific to the pole cells during the blastoderm stage and the migrating germ cells after gastrulation (Fig. 3C1-C3).

*Nasvi2EG001470* encodes a short peptide of 80 amino acids and was not included in the most recent annotation of the *Nasonia* genome at NCBI, but was present in OGS 2.0 [49]. A very similar sequence is annotated in the close relative *Trichomalopsis* (JAL personal observation), indicating that it is a *bona fide* transcript that is either novel, or very rapidly evolving. *Nasvi2EG001470* is strongly expressed in the oosome and pole cells, while levels markedly decrease in migrating germ cells (Fig. 3D1-D3).

Besides the expression in the oosome and the pole cells, *Nv-endoglucanase* is initially expressed at both poles during pre-blastoderm stage and early blastoderm stage (Fig. 3E1-E2). Later in blastoderm stage, the expression is down-regulated at the anterior pole and become specific to the pole cells (Fig. 3E3). Proteins of this type are found extensively in Hemimetabola, Hymenoptera, and Coleoptera, and appear to have been lost at the origin of Lepidoptera and Diptera clades. It is intriguing that what roles this protein plays during *Nasonia* embryogenesis.

Five transcripts localized to the oosome enter and are maintained in the pole cells but are then downregulated in the migrating germ cells (Fig. 3F1-J3). This set includes homologs of *Drosophila anillin* (*scraps*), *qin, and innexin1 (ogre),* (*Nv-anillin, Nv-qin*, and *Nv-innexin1*, respectively). Anillin is an actin binding protein that localizes to the contractile ring during cytokinesis [50]. In *Drosophila* Anillin protein is localized in the cleavage furrows when forming the PGCs [51], but its mRNA is ubiquitous. The early oosome localization of *Nv-anillin* mRNA suggests that the protein might also play roles in oosome outside of a potential conserved role in pole cell formation.

*Nv-qin* encodes a protein containing tudor domains along with an E3 ubiquitin ligase domain. *qin* is important in processing germline piRNAs, repressing retroelements assembling the nuage, and proper completion of oogenesis in the fly [52-54]. While *qin* has an important late role in germline cells, it is only weakly and diffusely expressed during embryogenesis in *Drosophila* [55]. Its mammalian homolog *RNF17* is required for production of specific particles in the germline nuage and for sperm development, but not for early germline specification [56].

*Nv-innexin1* encodes a putative gap junction protein whose fly homolog is most well-known for its role in proper development and function of the nervous system [57]. Other unexpected roles for innexin proteins have been described and proposed in insects [58], but at the moment the potential functional significance of the germline localization in *Nasonia* is unclear.

Two transcripts localized to the oosome and preserved in the pole cells have not clear *Drosophila* homologs (Fig. 3I1-J3). One of these (*Nv-C16orf87*) encodes a homolog of the human protein *C16orf87* and is expressed in the posterior region as well as specifically in the oosome in pre-blastoderm stage (Fig. 3I1-I3). The protein of *Nv-C16orf87* belongs to the uncharacterized protein family UPF0547, which contains the zinc-ribbon motif, and functions of this protein and it homologs are not known. It appears that this gene has been lost specifically in the Brachyceran fly lineage as it is found in beetles, moths, and some mosquitos, but not *Drosophila* (JAL personal observation).

Finally, another ankyrin domain encoding transcript is strongly localized to the oosome and is taken up into pole cells (Fig. 3J1-J2). It later has a complex and dynamic pattern in the blastoderm stages (Fig. 3J2-J3, additional images not shown). This transcript is a member of the newly described CLANK (Chalcid Lineage-specific ANKyrin-domain gene) family, of which there are nearly 200 in the *Nasonia* genome [43]. To differentiate from the others, we name it *Nasonia vitiripennis Oosome CLANK* (*Nv-OoCLANK*).

### Transcripts enriched in the oosome but excluded from pole cells

A set of 12 genes is expressed in the oosome but not transported to the pole cells. We predicted that these would have germline roles primarily in the oosome itself, or in the early stages of pole cell formation. We also expect many transcripts in this set will have roles outside of germline production, such as in embryonic patterning (as already known for *Nv-dpp* and *Nv-cad*) [59, 60]. Examples of what we consider potential embryonic patterning factors include: a CLIP protease encoding message (*Nv-mp1*) related to fly Melanization Protease and Easter (Fig. 4A1-A3), *Nv-kayak* (Fig. 4 B1-B3) encoding at transcription factor downstream of JNK signaling[61], and *Nv-elbow* (Fig. 4C1-C3) encoding a single zinc-finger transcription factor [62]. *Nv-kayak* is later expressed in a dorsal domain, indicating a conserved role in extraembryonic patterning. Interestingly, *Drosophila elbow* interacts with *orthodenticle* in specifying the ocelli and in photoreceptor cell fate determination [63], and an intriguing possibility is that these proteins work together in posterior patterning, with *Nv-elbow* possibly playing an important role in differentiating the posterior targets of *otd* from the anterior ones.

Several oosome resident transcripts have suggestive functional annotations. For example, the *Nasonia coronin* gene (*Nv-coronin,* (Fig. 4D1-D3)) encodes a protein whose homologs are known to bind and modulate actin, provide links between the actin and microtubule cytoskeletons, and regulate endo- and exocytosis in several developmental contexts [64, 65]. A germline role for the *Drosophila* Coronin ortholog has not been observed. *Nasonia milton* (*Nv-milton*) is another exciting transcript (Fig. 4E1-E3). *Drosophila* Milton is an adaptor protein that allows mitochondria to be loaded onto, and transported by, microtubule motors [66].

An oosome resident mRNA encodes a protein whose fly ortholog is uncharacterized, but whose function may be relevant to oosome function. This is the *Nasonia* homolog of *CG4552* (*Nv-CG4552*), which encodes a protein with a TBC25 domain (Fig. 4F1-F3). Proteins with this domain interact with Rabs to regulate membrane trafficking and dynamics. Such activities have been shown to be crucial for Osk function in the fly [67], and *Nv-CG4552* may play a supporting role in regulating membrane dynamics in the wasp. Another suggestive localized factor does not have clear orthologs outside of the hymenoptera, but it does have two predicted RNA Recognition Motifs, therefore we name it *Nv-rrm* (Fig. 4G1-G3). RRM domains bind RNA and are components of proteins that regulate RNA localization and translation. This novel lineage specific protein could therefore be involved in the localization of specific RNAs in the oosome, or the regulation of translation of specific RNAs within it.

Many of the oosome localized transcripts do not have annotations that lead to simple hypotheses about their roles in specifying the germline. One of these is *Nv-waterproof*, which encodes a fatty acyl-CoA reductase. *Drosophila waterproof* produces the hydrophobic molecules that coat the trachaeal tubes during *Drosophila* embryogenesis and is essential for gas filling of the trachea [68]. The protein’s novel role in *Nasonia* germline is not clear and worth to investigate, as is its early maternal expression in the oosome and later zygotic expression as the dorsal strip and in the extraembryonic tissue. (Fig. 4H1-H3). *Nv-CG42269* encodes a predicted organic ion transporter protein whose *Drosophila* homolog (*CG42269*) has no described function (Fig. 4 I1-I3).

Three oosome localized transcripts have no clear homologs in *Drosophila* or in other model organisms. *LOC100123551* has a sterile alpha motif (SAM) domain, which might indicate protein-protein or -RNA interactions (Fig. 4J1-J3). *LOC103315681* contains weak similarity to the N-terminal domain of Folded-gastrulation proteins (but is not a folded gastrulation ortholog) (Fig. 4 K1-K3), while *Nasvi2EG022821* has no discernible conserved domains (Fig. 4L1-L3). The functions of these factors will be the object of future investigation.

### Transcripts enriched in the posterior pole but not specifically the oosome

There are five transcripts that are significantly enriched in the posterior region of the embryo, but do not show significant association with the oosome (Fig. 5). The early embryonic expression for these transcripts appears as a cap or broad posterior to anterior gradient of mRNA. The significance of such transcripts to oosome assembly or to germ cell formation is not clear. Two of this class are known developmental transcription factors: orthologs of Zerknuellt [69], and Mothers against dpp [70] (*Nv-zen* and *Nv-mad2,* respectively (there are two mad paralogs in *Nasonia*). Two transcripts are predicted to encode catalytic enzymes: a choline kinase homologous to the *Drosophila CG2201* (*Nv-CG2201*), and a homolog of the ADP ribosylation factor-like 4 protein (*Nv-ARL4*). Finally, *LOC100680197* and *LOC100677932* have no identifiable homologs outside of hymenoptera. *LOC100680197* encodes a protein with MYND-type zinc-fingers and a p27-like domain, while *LOC100677932* has no clear conserved or functional domains.

### Functional analysis by parental RNA interference showed low phenotypic penetrance

While localization of an mRNA to the oosome and pole cells may strongly suggest a function related to PGC specification, demonstration of any such function is required. We chose a sampling of five promising molecules for in depth functional analysis (*Nv-bark, Nv-anillin, Nv-rrm, Nv-coronin*, and *Nv-innexin1*).

We initially tried to apply our parental RNAi (pRNAi) approach [44], but quickly found that this was not the ideal approach. Most dsRNAs caused partial sterility, with most of the obtainable eggs being apparently normal escapers. Eventually, we managed to collect embryos with phenotypes in a very low penetrance (2%-6%) from three of the five genes that we studied with the dsRNA concentration of 1ug/uL to 2.5ug/uL. These genes are *Nv-rrm, Nv-coronin*, and *Nv-innexin1*. They all either have no pole cells formed or less pole cells formed with and disorganized germ plasm residue at the posterior pole of the embryos in blastoderm stage. Infrequently, *Nv-coronin* knockdown embryos were characterized by pole cells that did not migrate to the gonad, but instead remained at the pole after gastrulation. (data not shown).

This issue was more serious for *Nv-bark* and *Nv-anillin*. At concentrations from 1.5ug/uL-2.5ug/uL), there were no eggs laid. At lower concentrations (250ng/uL, 500ng/uL and 750ng/uL), the eggs we collected all showed normal development. Since phenotypic embryos were either completely absent or extremely rare for our genes of interest, it became necessary to develop a new technique to assess the functions of the novel oosome genes we discovered.

### Development of an embryonic injection protocol for *Nasonia* RNAi

To circumvent the low penetrance problem from the pRNAi, we developed a protocol for embryonic injection of dsRNA followed by fixation and *in situ* hybridization (see details in Methods and Materials). As a negative control, we injected dsRNA against eGFP to test whether the physical injection and the dsRNA itself would affect the structure of the oosome and formation of the pole cells non-specifically. We were happy to find that even in embryos with obvious physical damage, the oosome and pole cells could form normally (Fig. 6A1-A4).

**Fig. 6.**
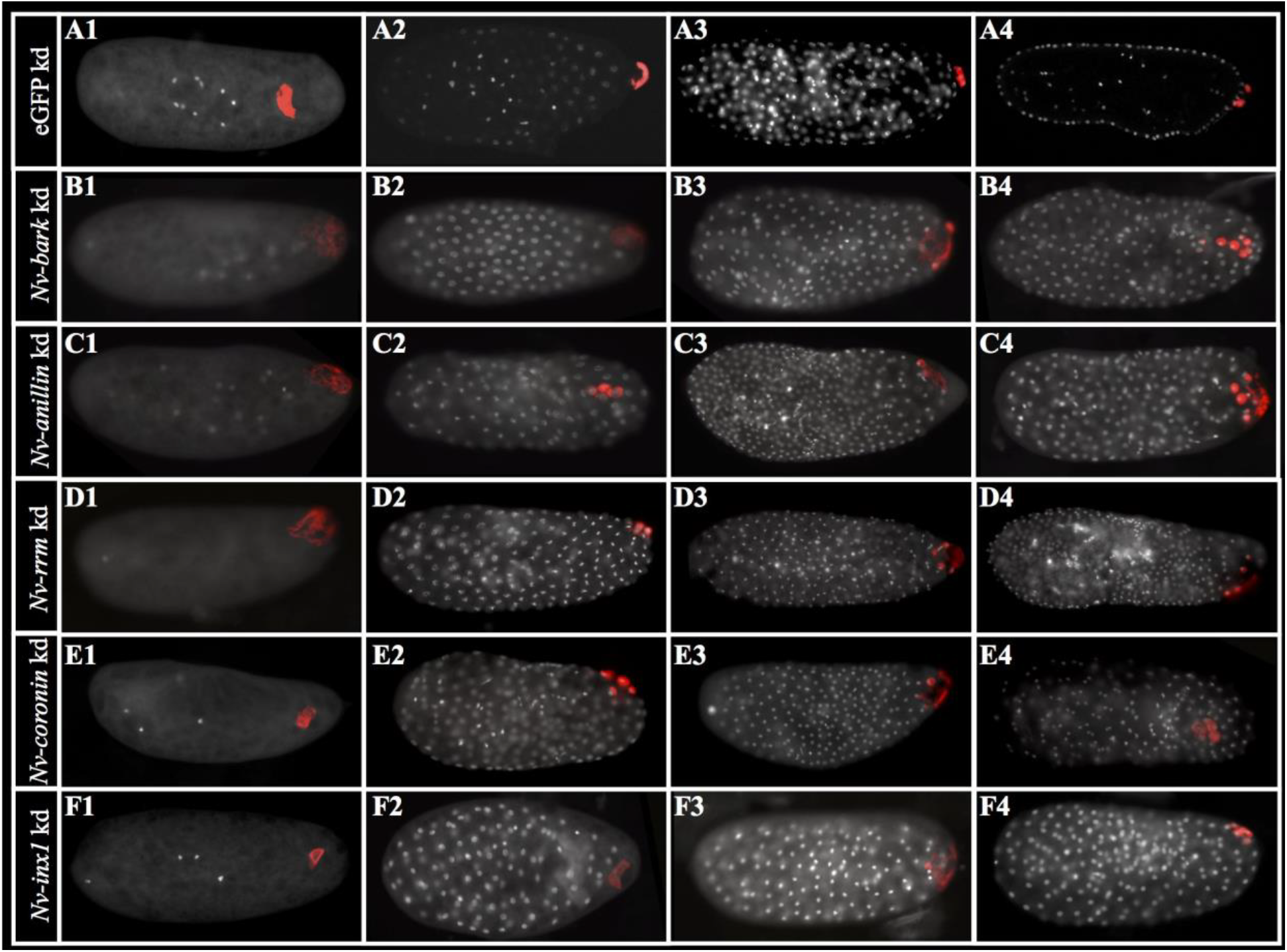
Functional analysis of a subset of germplasm localized factors by eRNAi. A1-A4: eGFP dsRNA injected embryos of increasing as the negative control. B1-B4: *Nv-bark* dsRNA injected embryos, C1-C4: *Nv-anillin* dsRNA injected embryos, D1-D4: *Nv-rrm* dsRNA injected embryos, E1-E4: *Nv-coronin* dsRNA injected embryos, F1-F4: *Nv-innexin1* dsRNA injected embryos. All the embryos are positioned with posterior side to the right and dorsal side on the top. The red in B1-B4 are marked by *Nv-nos*, and the rest are marked by *Nv-bark*. DAPI (white) marks the nuclei.

Of course, at a certain point, damage becomes too severe, leading to the death of the embryo. We set a stringent criterion for collecting embryos for later analysis by removing those where yolk leakage exceeded more than 10% of the embryo size (although all embryos with this amount of damage showed normal germ plasm and germline development) (Fig. 6A2, A4). We also excluded embryos that showed major morphological changes as compared to uninjected embryos.

After removing the embryos with obvious major damage both after injections, and when performing eggshell dissection after the fixation, we were left with about 85% embryos with viable embryogenesis by the time of imaging after *in situ* hybridization. The same criteria were also applied to the experimental knockdowns where the percentages were roughly the same as in the negative control When determining the phenotypes for the five genes, we considered the disruptions of developmental events to be potential effects of a knockdown when they were only specific to the knockdown and were never observed in the negative control.

We performed the same procedures when injecting and collecting the embryos as described for the negative control. The penetrance given by the embryonic RNAi (eRNAi) knockdowns is higher than the pRNAi knockdown, ranging from 20% to 39% across experiments. Phenotypes are evident in both the pre-blastoderm and pole cell stages, for *Nv-bark, Nv-anillin* and *Nv-rrm*. In contrast, the oosome is not affected in pre-blastoderm embryos after *Nv-coronin* and *Nv-innexin1* dsRNA injection, with phenotypes becoming evident only after the pole cells should have formed.

### RNAi against three novel germ plasm components unexpectedly disrupts the oosome at from an early stage

One transcript that particularly captured our attention was *Nv-bark*, as it was the transcript most strongly and consistently associated with PGC specification over embryonic development. However, its potential function in the germline is not clear. Since it encodes a transmembrane protein involved in epithelial junctions in *Drosophila*, we speculated that it might have a role in mediating adhesion or migration of the pole cells once they were formed. Surprisingly, the phenotypes produced by knocking down this transcript showed a much earlier requirement for this transcript. In early embryos (before migration of nuclei to periphery), the oosome has lost its integrity as a single unit (Fig. 6B1). Instead there are scattered particles of what appears to be oosome-like material, concentrated around the lateral cortex of the embryo. Later, at the time the pole cells would normally form, no budding is observed, and some loose aggregates of germ plasm like material remains attached to the cortex (Fig. 6B2-B3). In some cases, germ plasm surrounds nuclei in a way similar to what is seen in pole cells (Fig. 6B3, B4). However, these nuclei remain part of the embryonic syncytium.

Like *bark,* the mRNA of *Drosophila anillin* is not localized to the polar granules [71]. Anillin protein, however, accumulates at the base of pole cells when they are budding in *Drosophia* [51]. Since Anillin is a major component of the contractile ring during mitotic cytokinesis [50] and is enriched at the bud furrow during pole cell formation in *Drosophila* [51], we predicted that the enrichment of *Nv-anillin* mRNA in the oosome and pole cells would be related to *Nasonia’s* unique way of forming a single large pole bud instead of several small ones, as occurs in *Drosophila*. Surprisingly, the phenotype of *Nv-anillin* is indistinguishable from that of *Nv-bark*: The oosome does not form properly, and germ plasm material remains bound to the posterior cortex of the embryo, leading to the association of the syncytial nuclei at random posterior locations (Fig. 6C1-C4). The same set of phenotypes is observed when we knock down *Nv-rrm* with eRNAi (Fig. 6D1-D4). Thus, three genes with very different predicted functions all result in the same phenotype.

It is important to note at this point that the common phenotype of the above three knockdowns is not the same as the complete loss of germ plasm activity. Such phenotypes are seen for *Nv-osk, Nv-vas*, and *Nv-aubergine (Nv-aub).* When these genes are knocked down, posterior mRNAs such as *Nv-nos* take on a uniformly graded posterior cap and no enriched accumulation of germ plasm markers is ever observed at the posterior [35, 40]. This indicates that these transcripts are involved in the specific form of the oosome, but may not be essential for the production of germline-like cells.

Finally, despite the fact that the oosome does not form or migrate through the embryo as in wildtype, a bud like protrusion similar to the one that initiates pole cell formation is regularly observed (Fig. 6C3, Fig. 1E), and this bud is not associated with the bulk of the germ plasm like material. This indicates that neither the oosome, nor its remnants, induce the bud, and that bud formation may be autonomous to the embryo. This has some precedent in *Drosophila*, where the autonomous ability of the fly embryo to produce pole-cell like structures at both poles of the embryo is revealed when Arf guanine exchange factor Steppke is reduced [72]. However, in the fly, normal global repression of pole-cell formation is overcome by germ plasm components (primarily *germ-cell-less*), while in *Nasonia,* at least the initial budding appears to be germ plasm independent.

### RNAi against *Nv-coronin* and *Nv-innexin1* does not affect the oosome, but disrupts pole cell formation

We tested the functions of two other transcripts with eRNAi: *Nv-coronin* and *Nv-innexin1*. Knockdown of both genes left the oosome intact, and able to migrate through the embryonic cytoplasm normally (Fig. 6E1, F1). However, in both cases, pole cell formation fails, indicating that these genes have downstream functions that are specific to pole cell formation (Fig. 6E2-E4, F2-F4). *Nv-coronin*’s functional annotation is consistent with a role in cellularization, as it is predicted to interact with both the microtubule and actin cytoskeletons, both of which are crucial for mitosis and cell formation. While Innexin1 is most well known as a crucial component of gap junctions and a role in the nervous system [57], a role for at least one Innexin in cellurization has been demonstrated in the beetle *Tribolium* [58], suggesting that the full potential of these proteins in regulating cellular processes has not been fully explored.

## Discussion

### RNA-seq analyses

Our results have uncovered an unexpectedly large amount novelty in the mRNA content of the germline determinant of the wasp *Nasonia vitripennis*. This was achieved using RNAseq followed by statistical detection of differential expression of mRNAs between the anterior and posterior poles of the early embryo. The statistical predictions were then extended by *in situ* hybridization and RNAi of candidate genes.

Our goal was to identify all transcripts specifically localized to the oosome, and it is worth contemplating how close we came to achieve this goal. We used three different approaches that varied in the strategy for mapping, quantifying transcripts and assessing differential expression. These analyses agreed on the vast majority of the genes with putative significantly different enrichment at the two poles, and most of the disagreement was at the margin of differential expression that would be likely biologically significant. In addition, all analyses found genes previously identified to be localized anteriorly or posteriorly. We even detected all known genes with both strong oosome localization and strong localization at the anterior pole. We thought these could be missed, because if the anteriorly localized mRNA population approached the same levels as that in the oosome, the enrichment might be obscured.

For these reasons, we believe that we have uncovered the vast majority of mRNAs localized to the *Nasonia* oosome. Of course, no approach can guarantee comprehensiveness, and some molecules may have been missed for several reasons. For example, while the *Nasonia* genome is well annotated, it is possible that a very few genes are not represented in the genome assembly or the transcriptome annotations, and thus would not have been assessed. Indeed, we did identify slightly different sets of genes when using NCBI annotation 102 versus the OGS 2.0 annotation [49], which were created using different approaches to identify and predict genes. Most of the differences are at the margins of significance and low levels of differential expression. However, there are a handful of confirmed localized transcripts detected using annotation 102 that were missed in analyses with OGS 2.0. In addition, at least one transcript was not annotated in annotation 2.1, but was found in annotation 2.0 and was confirmed to be localized to the oosome (*Nasvi2EG001470*, Fig 3D). This is likely a false negative annotation in annotation 102, as a very similar sequence is found in the transcriptome of the very closely related wasp *Trichomalopsis sarcophagae* [73].

Another issue that can cause false negatives is the large number of *Nasonia* transcripts that overlap, which can lead to the concatenation of transcripts. If a localized transcript is fused to a ubiquitous, highly expressed transcript, the signal of localization can be lost. This problem is largely solved by preventing novel junctions when mapping with tophat2. However, this also seem to change the calculation of significance for some transcripts, so performing the analysis with and without novel junctions gives more complete results.

Finally, there are some unknown artifacts which may cause significantly enriched genes to be missed. A prime example of this is our discovery of *Nv-coronin*. A preliminary attempt at this experiment resulted in generally poor sequencing results, and a completely unusable replicate. In general, this analysis gave predictably worse results, where several known localized genes were not found to be significant. However, all of the molecules whose functions were analyzed in this manuscript were found to be statistically significantly enriched at the posterior, including *Nv-coronin*, and we proceeded to clone them to test their localization and function. In the subsequent analyses based on high quality sequencing results (used as the basis for this manuscript), *Nv-coronin* was excluded by cuffdiff for statistical testing for unknown reasons, despite showing similar posterior enrichment that had previously been deemed statistically significant. Thus, there is a potential for false negatives if this artifact affects several genes. Very few transcripts in the “not tested” category show a similarly strong posterior enrichment, so we believe that this effect is also small.

### Comparison of the oosome of *Nasonia* to the polar granules of *Drosophila*

Germ plasm in insect embryos must perform several different functions. First, it must be able to concentrate and arrange the set of proteins and RNAs (e.g. Tud, Vas, *nos* mRNA, Piwi/Aub) that are associated with germline fate in the posterior of the egg. Second, it must ensure its own incorporation into the PGCs as they form by interacting with the specialized cytoskeletal structures that mediate pole cell formation. Germ plasm must also include molecules involved specialized features of PGCs. Such functions include the concentration and selection of mitochondria, repression of transcription, repression of transposons, and guidance of the germ cell migration to the gonad primordia.

On the other hand, we had already observed that the oosome’s morphology and interaction with the cytoskeleton (during its migration and formation of the pole cells) was quite distinct from *Drosophila*, as is the way the pole cells migrate into the interior of the embryo. Therefore, we expected to find a mixture of conservation and novelty when we examined the mRNA content of the oosome. Indeed, this is what was found, but with a surprisingly strong bias toward novelty.

In terms of conservation, our analyses found *Nv-nos, Nv-osk* (already known factors found in both oosome and polar granules) as well as *Nv-ovo*. We also found *Nv-aub* whose ortholog is localized as protein, but not mRNA, in *Drosophila*. While there are a large number of polar granule localized mRNAs that are not found in the oosome, we will only discuss a few that are significant.

mRNA for *polar granule component (pgc)* encodes a small peptide [74], and is strongly localized to the posterior pole [75]. Polar-granule-component protein has a crucial role in the global repression of transcription that occurs in pole cells upon their formation, through an interaction with the transcription elongation factor TEF-b [74]. This repression is a widely conserved feature of PGCs across animals, which makes it somewhat surprising that *pgc* appears to be a novelty in the *Drosophila* lineage [75].

Fascinatingly, TEF-b may be a unifying factor underlying germline quiescence, as it interacts with PIE-1 in the worm *C. elegans* to repress transcription [76]. In line with this, we have found a highly conserved transcription elongation factor, *Nv-spt5* (Fig. 3 A1-A3) localized strongly to the oosome and pole cells. In human cells, Spt5 acts as an inhibitor or transcriptional elongation, until its C-terminal domain is phosphorylated in a TEF-b dependent manner [77]. If an interaction between *Nasonia* Spt5 and TEF-b does have a role in regulating the cessation of transcription in *Nasonia* pole cells, it would be strong evidence for TEF-b being a core conserved component of the germline fate, whose interaction partners and regulators are labile across lineages.

Another crucial *Drosophila* germ cell factor that is not present in the *Nasonia* oosome is *germ cell less* (*gcl*). The Germ-cell-less protein itself is very highly conserved at the sequence level in *Nasonia,* but the mRNA showed no enrichment in our RNAseq experiments, and we independently confirmed by *in situ* hybridization that it is expressed uniformly throughout the early embryo (not shown). Gcl is important for the proper production of pole cells, apparently by regulating the orientation centrosome separation at the posterior pole, which is required for efficient pole cell formation and uptake polar granules by the pole cells [78]. At the molecular level, Gcl seems to act by downregulating torso signaling, to allow the proper conditions for pole cells to form. The lack of Gcl function in the germline of *Nasonia* is consistent with the lack of Torso signaling at the termini in the wasp [79], making the need for Gcl redundant. At the moment, it is not clear whether the use of Gcl in pole cell formation is a recent novelty in *Drosophila*, or whether it was present ancestrally, but lost in the Hymenopteran lineage.

A number of transcripts found in the *Nasonia* oosome are good candidates for generating essential PGC features. For example, a high concentration of mitochondria is a strongly conserved feature of germ plasm and PGCs across animals [80, 81]. In *Drosophila*, the long Osk isoform plays an important role in concentrating mitochondria in the pole plasm [82]. But, since the long form of Osk appears to be a novelty of *Drosophila* and its close relatives, other molecules should be expected to perform this role in other species. Suggestively, mRNA encoding a Milton ortholog was found strongly localized to the *Nasonia* oosome. Milton acts an adaptor that loads mitochondria onto microtubule motors for transport and localization within and between cells in *Drosophila* [83], and we propose that *Nasonia* Milton plays a role in enriching mitochondria around the oosome and in the pole cells in the wasp, and perhaps other insect species that lack the specialization of long Osk isoform.

Another critical function for germ cells is the control of transposable elements, which is often dependent on Tudor domain containing proteins. mRNAs for two Tudor domain proteins are present in the oosome, including *Nv-qin* and *Nv-tdrd7*. Neither of them is enriched in the polar granules or pole cells in *Drosophila*, but both have crucial roles during oogenesis to reduce the activity of transposable elements [52, 53, 84]. The presence of these additional Tudor domain encoding transcripts may indicate that either there is an increased activity of transposable elements in *Nasonia* that requires an earlier response, or perhaps other mechanisms are employed in *Drosophila* to combat transposon activity in the early PGCs. Further sampling of germ plasm of other insects should help to resolve these questions.

Germ cells are known to have a distinct metabolic profile from somatic cells, and this difference is related to their pluripotent stem cell-like properties, and to the requirements of their migratory properties [85, 86].Potentially related to this we have found that a *Nasonia* insulin-like growth factor I mRNA is localized to the oosome. Interestingly, this mRNA encodes a short protein that shows much stronger similarity to insulin proteins of vertebrates, than it does to any of the insulin-like molecules of *Drosophila* (DILPs) (JAL personal observation). In addition, an mRNA encoding a putative organic cation transporter (*Nv-CG42269)* containing a Major Facilitator Superfamily (MFS) domain is strongly localized to the oosome. Such molecules are crucial for regulating cellular metabolism and signaling at multiple levels, by controlling the trafficking of many small organic molecules (including sugars) within and between cells [87, 88]. Lipid metabolism is also uniquely regulated in germ cells, and the identification of the *Nasonia* homolog of the Acyl-CoA reductase *waterproof* may reflect this [89]. Finally, we surprisingly found a transcript encoding a protein similar to endoglucanases found in several insect lineages (but absent from Diptera). Enzymes of this type are broadly defined by their ability to break down polymers glucose. The substrate and potential role for this enzyme in the germline is not yet known.

In addition to providing insight into the conserved functions of germ cell components, we also found several molecules that do not have clear homology outside of the Hymenoptera, or in some cases outside of *Nasonia* and its closest relatives. This includes a novel RNA recognition domain containing protein whose function we analyzed in depth (*Nv-rrm* discussed in the following section). We also found that an mRNA encoding an ankyrin domain protein (*Nv-OoCLANK* (*LOC100679945*)) that belongs to a family of proteins that underwent a massive amplification within chalcid wasp lineage, and which appears to have entered the ancestral chalcid wasp by horizontal transfer [43]. Finally, a handful of transcripts have no identifiable domains or homologs. The functional relevance of these molecules will be an area of intense interest in the future.

### Unexpected functions of novel oosome components

Early in our analysis we chose a handful of transcripts for functional analysis. These were chosen based on a combination of criteria that included high enrichment in the oosome, novelty, and the potential to give phenotypes that we could characterize with the current set of functional tools available to us in *Nasonia*. Three of these (*Nv-bark, Nv-anillin* and *Nv-rrm*) gave an unexpected phenotype, where oosome-like material was not coalesced into the typical spherical oosome structure, but rather was scattered in clumps attached to the plasma membrane near the posterior pole. Eventually, these clumps of germ plasm-like material come into contact with syncytial nuclei when they migrate to the cortex. However, this remnant material is unable to induce the pole cell fate. It is important to reiterate that this phenotype is quite distinct from that seen for genes that have a core role in oosome assembly (*Nv-osk, Nv-tud, Nv-vas* and *Nv-aub* (not shown). In these cases, there is no hint of the oosome, and mRNAs normally localized in the oosome are distributed in homogenous caps at the posterior pole, rather than as discrete clumps of material [35].

These phenotypes indicate these genes are involved in the coalescence of the oosome into single entity within the central column of cytoplasm in the embryo, and/or maintenance of oosome integrity. The molecular bases of such functions are not completely clear at the moment for any of these three genes. *Nasonia* Bark is a putative transmembrane protein and would be predicted to be targeted to the membrane. One potential hypothesis is that *Nasonia* Bark is targeted to the membrane at the posterior pole of the embryo, releases and/or repels oosome material from adhering to the plasma membrane, forcing the oosome to remain in the bulk cytoplasm where it concentrated into a large sphere and moved around by strong cytoplasmic flows that occur during the earliest cleavages. This model would also imply that interaction with the cortex prevents oosome material from coalescing, leading to the scattered clumps we observe in *Nv-bark* knockdowns. Mechanistically, this could be related to the ability of Bark to induce endocytosis [90], a process that is associated with proper anchoring of the germ plasm and the recruitment of specialized actin binding proteins to the posterior pole in *Drosophila* [91].

An alternative hypothesis is that *Nasonia* Bark produced in the oosome is not secreted but is instead incorporated as an important structural component of the large, solid form of the oosome. This is consistent with the structure of the protein, which contains main protein-protein interaction domains of different types [45, 46]. Testing these hypotheses will require in depth analysis of the subcellular localization of *Nasonia* Bark during early embryogenesis, and proteomic analysis of binding partners of *Nasonia* Bark.

The role of *Nv-anillin* in maintaining the stability of the oosome was also surprising and the molecular basis of the phenotype will require further investigation. Anillin orthologs are well known as actin binding proteins involved in assembling the contractile ring required to separate cells in cytokinesis [50]. Anillin also plays a crucial and novel role in the specialized cytokinesis of the *Drosophila* pole cells [51]. While these known functions might have indicated that oosome localization of *Nv-anillin* was related to an important role in the specialized polar bud formed in *Nasonia*, RNAi showed that this protein has an earlier role in oosome assembly/maintenance (Fig. 6 C1). Similar to *Nasonia* Bark, one possible function of *Nasonia* Anillin is as a structural component of the oosome, which may or may not be related to its ability to bind actin and associated proteins. Alternatively, *Nv-anillin* may act to release and/or repel the oosome from the cortex, as proposed above for *Nv-bark*.

Anillin homologs have known functions which are directly related to the processes of release of germ plasm from the embryonic plasma membrane. In *Drosophila*, the germ plasm is tightly bound to the plasma membrane until the nuclei reach the posterior pole of the embryo. The centrosomes associated with these nuclei mediate detachment of the pole plasm from the cortex through interactions of the astral microtubules emanating from the centrosomes [92]. This is in contrast to *Nasonia,* where the oosome detaches from the cortex at about the same time as the zygotic nucleus begins its first division at the anterior pole. While nuclei are lacking at this time, numerous centrosomes are present, as they are provided maternally in a process characteristic of many Hymenopteran embryos [93]. *Nasonia* Anillin could be relevant to a model where astral microtubules emanating from maternally provided centrosomes detach oosome material from the cortex, because Anillin homologs have been shown to mediate interactions between the actin cytoskeleton and cortical and subcortical microtubule arrays in multiple model systems [94, 95]. Again, in depth examination of the subcellular localization and interactions of *Nasonia* Anillin will be required to completely understand its role in maintaining the oosome.

Based on the presence of only RNA recognition motifs in the protein, we predict that *Nv-rrm* will have one of two likely roles. One possibility is that it is involved in translational regulation of key regulators of oosome structure, presumably including *Nasonia* Anillin and *Nasonia* Bark. Alternatively, *Nv-rrm* may be important in binding RNA and protein in order to maintain the structural integrity of the oosome. In addition, neither of these possibilities is mutually exclusive. The knockdowns of *Nv-coronin* and *Nv-innexin1* had specific effects only on the formation of pole cells, while the oosome appeared to remain intact. Coronin is an actin binding protein associated with the formation of highly concentrated networks of F-actin [65]. It seems likely that *Nasonia* Coronin has an important role in organizing an actin cytoskeleton arrangement specialized for the formation of the large polar bud that initiates pole cell formation.

The potential role of *Nv-innexin1* is somewhat more mysterious. Innexins are typically known as components of the gap junctions that are found in some tightly integrated epithelial tissues. Such junctions would not be expected of the motile pole cells. Interestingly, Innexin-7 (a paralogous protein with a similar structure to *Nasonia* Innexin1) in the beetle *Tribolium* has a novel role in cellularization of the syncytial blastoderm. Such a function for *Nasonia* Innexin1 would explain the failure in pole cell formation we see after RNAi.

## Conclusion

This work has revealed numerous unexpected mRNAs that are localized to the germ plasm of the wasp *Nasonia.* The results have given insights on the potentially ancestral mechanisms used by germ plasm to accomplish conserved required functions, such as a possible ancestral role for Milton orthologs to bring mitochondria to the germ plasm, a function replaced by the long Osk isoform in *Drosophila.* On the other hand, our results have identified numerous components that are likely to be specific to *Nasonia* and its relatives in the parasitic wasp lineage (e.g., *Nv-OoCLANK, Nv-rrm*, and the use of Nv-Bark and Nv-Anillin in assembling the oosome). Deeper analyses of the functions of these molecules in *Nasonia*, and broader sampling of germ plasm in other holometabolous insects will be required to determine the patterns of evolutionary change in the germ plasm. Such analyses will be important because the germ plasm is a uniquely powerful organelle that can rapidly drive naive nuclei into a highly specialized, yet functionally totipotent state. Understanding how and why such a fundamental substance changes and is even lost in the course of evolution will provide foundational insights into the mechanisms of cell fate determination and the interaction of subcellular organelles and their cellular milieu.

## Methods

### Sample preparation

In order to identify the components of the maternally deposited mRNAs in the oosome located in the posterior half of the parasitoid wasp *Nasonia vitripennis* embryos, we collected and bisected the pre-blastoderm stage embryos (0-2 hours after egg lay at 25 °C) to detect the differential expression levels between the anterior and posterior halves of the embryos.

The embryos were aligned with the anterior pole to the right on the ice-prechilled and heptane glue-coated slide. Then a thin layer of halocarbon oil 700 (Sigma) was applied to cover the embryos. The slide was transferred on the dry ice-prechilled “guillotine” and then was anchored by tightening the screws on each end. After the halocarbon oil 700 was solidified on dry ice, put the guillotine on the dry ice-prechilled stabilizer and transferred it under the dissection microscope. The embryos were positioned to match the slot in the guillotine where the dry ice-prechilled razor blade will be inserted into. After the embryos were cut, the anterior and posterior halves of the embryos were immediately collected and transferred into the two 1.5mL non-stick RNase-Free microfuge tubes (Ambion) with the dry ice-prechilled probe, separately. Three biological replicates were created.

Total RNA was isolated from these six samples for library preparation. In the library preparation upon which this manuscript is based, we used around one microgram of total RNA from each sample, in which 100ng of total RNA was from the *Nasonia* other 900ug was from a distantly related parasitic wasp (*Melittobia digitata* [43]). Libraries were prepared using the NEBNext Ultra Directional RNA Library Prep Kit for Illumina (NEB #E7420) in conjunction with NEBNext Poly(A) mRNA Magnetic Isolation Module (NEB #E7490). Libraries were validated and quantified before being pooled and sequenced on an Illumina HiSeq 2000 sequencer with a 100 bp paired-end protocol. Sequence files are available in the NCBI SRA database under accession SRP156232.

### RNA sequencing data analysis

The quality of the sequencing data was determined using FastQC software. The sequences were processed by Cufflinks package for differential expression detection, using multiple variations on the default parameters (job files in Supplemental File 2). Briefly, raw reads were aligned to annotation 102 of assembly 2.1 of the *N. vitripennis* genom (https://www.ncbi.nlm.nih.gov/genome/annotation_euk/Nasonia_vitripennis/102/) using TopHat2[96]. These results were either used directly in cuffdiff [97], or were further processed using stringtie [98] to generate new transcriptome predictions and quantification. Various normalization parameters were used, and each permutation gave slightly different results. In addition, we mapped the reads using assembly 1.0 and annotation 2.0 (OGS 2.0) [49]. All of the commands jobs in this process of analysis are included in the Supplemental File 2. The cuffdiff results of all permutations of the analysis are provided in Supplemental File 3. The computing work was done by the High Performance Computing Cluster located at University of Illinois at Chicago.

Probes and dsRNAs for the chosen genes were generated by the protocol described in [40]. Primers for generating these templates are provided in Supplemental File 4. Alkaline phosphatase *in situ* hybridization was performed by the protocol described in [99].

### Embryonic RNA interference (eRNAi)

In order to perform eRNAi on the early *Nasonia* embryos to study the germline candidate genes’ functions, we created the following workflow:

Around 30 pre-blastoderm stage embryos (0-1 hour after egg lay at 25 °C) were collected and quickly aligned vertically on the heptane glue-coated 18mm×18mm coverslip. This coverslip was transferred and anchored on the ice-prechilled slide by applying a thin layer of water. The slide was then put in an air tight petri dish with proper amount of desiccant (Drierite with indicator, 8 mesh, ACROS Organics) pre-chilled at 4 °C. The embryos were dehydrated in the desiccant at 4 °C for 45 minutes. After dehydration, the embryos were covered with a layer of halocarbon oil 700 and were ready for microinjection.

The dsRNAs were dissolved in Nuclease-Free Water (Ambion) at the concentration of 1mg/mL and loaded into the Femtotips II Microinjection Capillary (Cat. No. 930000043, Eppendorf). The constant pressure was set at 500hpa and the injection pressure was set initially at 250hpa with periodic adjustment as the needle changed over the course of injection. The process of injection was performed at room temperature and needed to be done as soon as possible. After injection, the slide was transferred into a paper towel-moisturized petri dish pre-warmed at 28 °C to incubate the injected embryos for specific developmental stages. The embryogenesis of these embryos was stopped at pre-blastoderm stage (before the budding), beginning of blastoderm stage (during budding), and later in blastoderm stage (pole cells formed and/or after pole cell divisions). To stop the development, the coverslip was put into the heptane to wash off the halocarbon oil 700 for three minutes, and then transferred into the 37% formaldehyde-saturated heptane for 2-5 hours fixation in the dark with the embryos facing up.

After fixation, the coverslip was carefully taken out of the fixative, and flipped upside down to gently press the embryos on a double-sided tape that was taped on a petri dish, so that all the embryos can be anchored on the tape for dissection. Add about 15mL PBS with 1% Tween, use the needle (BD PrecisionGlide Needle, 30G × 1) to carefully remove the eggshells from the embryos. The dissected embryos were then transferred by pipette into the 1.5mL non-stick RNase-Free microfuge tubes. The embryos were immediately dehydrated by 100% Methanol and stored at −20 °C.

Before performing fluorescent *in situ* hybridization (FISH) on those eRNAi knocked out embryos, they need to be rehydrated with a series of Methanol/PBT washes (75%, 50%, 25%). The protocol for FISH was adapted from [40]. A detailed protocol is available on request.

## Declarations

### Ethics approval and consent to participate

Not applicable

### Consent for publication

Not applicable

### Availability of data and material

Sequencing results can be found in the NCBI SRA databaseat: **<u>Error! Hyperlink reference not valid.</u>**

### Authors’ contributions

JAL and HHQ collaborated on experimental design, analyzed the data and wrote the paper. HHQ performed the experiments and collected the data.

## Acknowledgements

We thank Daniel Pers for performing the combined *Nasonia/ Melittobia* RNAseq libraries, and the UIC RRC Sequencing core for arranging the sequencing of our samples.

## Additional Files

**Additional File 1. (.xlsx)** Significantly enriched transcripts from both experiments. The transcripts listed in “Compilation of Transcripts significant in the main analyses” were generated using the sequencing data from the second experiment by NCBI annotation 102 with various normalization parameters. The “Transcripts from the second experiment only found in OGS 2.0” were the transcripts from the second experiment only found using the annotation OGS 2.0, butnot found using NCBI annotation 102. The “Transcripts from the first experiment” were the transcripts only found in the first time sequencing data, but not in the second time sequencing data.

**Additional File 2. (.docx)** Command jobs. The command jobs for analyses using the NCBI annotation 102 and the annotation OGS 2.0 with various normalization parameters.

**Additional File 3. (.xlsx)** Original lists of the transcripts generated by Cuffdiff. This file includes the original lists of the transcripts generated by Cuffdiff using the NCBI annotation 102 and the annotation OGS 2.0 with various normalization parameters.

**Additional File 4. (.xlsx)** Primer list. All primers used in this manuscript are listed. This list includes the primers used for cloning the genes and making templates for the probes and dsRNAs.

